# Smad9 is a key player of follicular selection in goose via keeping the balance of LHR transcription

**DOI:** 10.1101/213546

**Authors:** Daolun Yu, Fanghui Chen, Li Zhang, Hejian Wang, Jie Chen, Zongmeng Zhang, Jie Li, Chaofeng Xing, Honglin Li, Jun Li, Yafei Cai

## Abstract

The egg production of poultry depends on follicular development and selection. However, the mechanism of selecting the priority of hierarchical follicles is completely unknown. Smad9 is one of the important transcription factors in BMP/Smads pathway and involved in goose follicular initiation. To explore its potential role in goose follicle hierarchy determination, we first blocked Smad9 expression using BMP typeⅠreceptor inhibitor LDN–193189 both *in vivo* and *in vitro*. Unexpectedly, LDN–193189 administration could dramatically suppress Smad9 level and elevate egg production (7.08 eggs / bird, *P*< 0.05) of animals, and the estradiol (E_2_) and luteinizing hormone receptor (LHR) level were significantly increased (*P*< 0.05), but the progesterone (P_4_) and follicle stimulating hormone receptor (FSHR) mRNA remain unchanged. Surprisingly, Smad9 knockdown notably attenuated (*P*< 0.05) in E_2_, P_4_, FSHR and LHR level in goose granulosa cells (gGCs). Further chromatin immunoprecipitation (ChIP) assay of gGCs revealed that Smad9, served as a sensor of balance, bound to the LHR promoter regulating its transcription. These findings demonstrated that Smad9 is differentially expressed in goose follicles, and acts as a key player in controlling goose follicular selection.

**SUMMARY STATEMENT:** To study the hierarchical development mechanism of avian follicle, new strategies can be found to improve the egg production of low-yielding poultry, such as geese.

## INTRODUCTION

The geese (*Anser cygnoides*) are cultivated widely in China as one of most important farm poultries. However, the poor fertility seriously restricts its further application. The avian ovary contains large number of follicles during the laying period, which can be divided into two types: prehierarchical follicles and preovulatory follicles. The prehierarchical follicles comprise three types of follicles: small white follicles (SWF, 1-2 mm), large white follicles (LWF, 3-5 mm) and small yellow follicles (SYF, 6-9 mm). The preovulatory follicles consist of yolk-filled follicles at five stages (F5, F4, F3, F2, F1, from small size to large size) in laying hens (Etches and Petitte, 1990; Jia et al., 2010). Folliculogenesis involves a series of events during which a growing follicle either develops to the ovulation stage or undergoes atresia (McGee and Hsueh, 2000). The maturation and ovulation of the follicle is strictly regulated by a variety of endogenous factors.

Bone morphogenetic proteins (BMPs) belong to the transforming growth factor β (TGF-β) superfamily and elicit their effects through activation of type-1 and type-2 serine/threonine kinase receptors on the target cells (Su et al., 2009). BMP type| receptor kinases phosphorylate various downstream receptor-regulated Smads (R-Smads) upon ligand binding (Kawabata et al., 1998; Schmierer and Hill, 2007). R-Smads include Smad1, Smad5 and Smad9 (also known as Smad8) can be phosphorylated by BMP type I receptor kinase form complexes with Smad4, a Co-Smad, and bind to the responsive element (BRE) to regulate the transcription of target genes (Massague et al., 2005). Therefore, R-Smads are vital for intracellular transmission of BMP signals.

BMP/Smads signaling pathways were reported to play important regulatory roles in the development, atresia and selection of the follicles in mammals (Costello et al., 2009; Haugen and Johnson, 2010; Knight and Glister, 2003; Miyoshi et al., 2007; Nilsson and Skinner, 2003; Pangas and Matzuk, 2004; Wang et al., 2010; Ying et al., 2001). However, there are clear differences in the follicular development between birds and mammals, and it is unclear whether the BMP/Smads system still plays a role in the folliculogenesis of birds. Smad9 is one important member of the Smads family proteins, although their function is largely unknown. Previous studies have shown that activation of BMP signaling pathway could increase Smad9 expression (Tsukamoto et al., 2014). Interestingly, Smad9 contains a special linker region that can result in its markedly lower transcriptional activity than Smad1 and Smad5. In turn, Smad9 overexpression can reduce the activity of BMP in a dominant negative manner through inhibiting the transcription of target genes (Tsukamoto et al., 2014). Therefore, Smad9 is crucial for the balance of BMP/Smads signaling. In addition, Smad9 was found to be involved in the follicular initiation but was silent during maturation process in goose (Xu et al., 2015). However, whether it is involved in regulating follicular development and the underlying mechanism is still unclear. To address these issues, Chinese Wanxi White Goose, which have a low annual egg yield of only about 20-25 eggs, were chosen in this study. The distribution of Smad9 in the ovary was investigated and then its expression was manipulated by using LDN-193189, a kinase inhibitor of BMP type Ⅰ receptors. Finally, we analyzed the hormone secretion, receptor-expression and the number of ovulation. Our results showed that Smad9 can regulate E_2_ secretion and LHR transcription both *in vivo* and *in vitro*. Most importantly, we found that attenuation of Smad9 expression during laying stage could significantly elevate goose egg production. This study shed new light on the role of Smad9 in avian follicular development and ovulation.

## RESULTS

### Smad9 expression in the GCs layer of follicles

To analyze the expression of Smad9 in goose ovary, the ovaries were collected from Chinese Wanxi White Goose in laying period. We can see a number of prehierarchical follicles and only one preovulatory follicle at different development stages in the mature goose ovary (Fig. 1A). Hematoxylin and eosin (H&E) staining was further carried out to analyze the histological structure of follicular wall. The results revealed that the thickness of GCs layer showed a trend of first increased and then decreased, which reached the maximum at F3. No significant difference in the thickness of GCs layer among SWF, F5 and F2 as well as difference between SYF and F1 was detected, however significant differences were shown among F3, F4, F1 and SWF (*P*< 0.05) (Fig. 1B). Finally, immunohistochemical analysis showed that Smad9 was specifically expressed in the GCs layer of follicles, and no signaling was detected in other parts of the ovary (Fig. 1C). Based on these observations, it can be concluded that the morphological of GCs layer showed regular changes with follicles development and the Smad9 was expressed only in GCs layer in goose follicles.

**Fig. 1.**
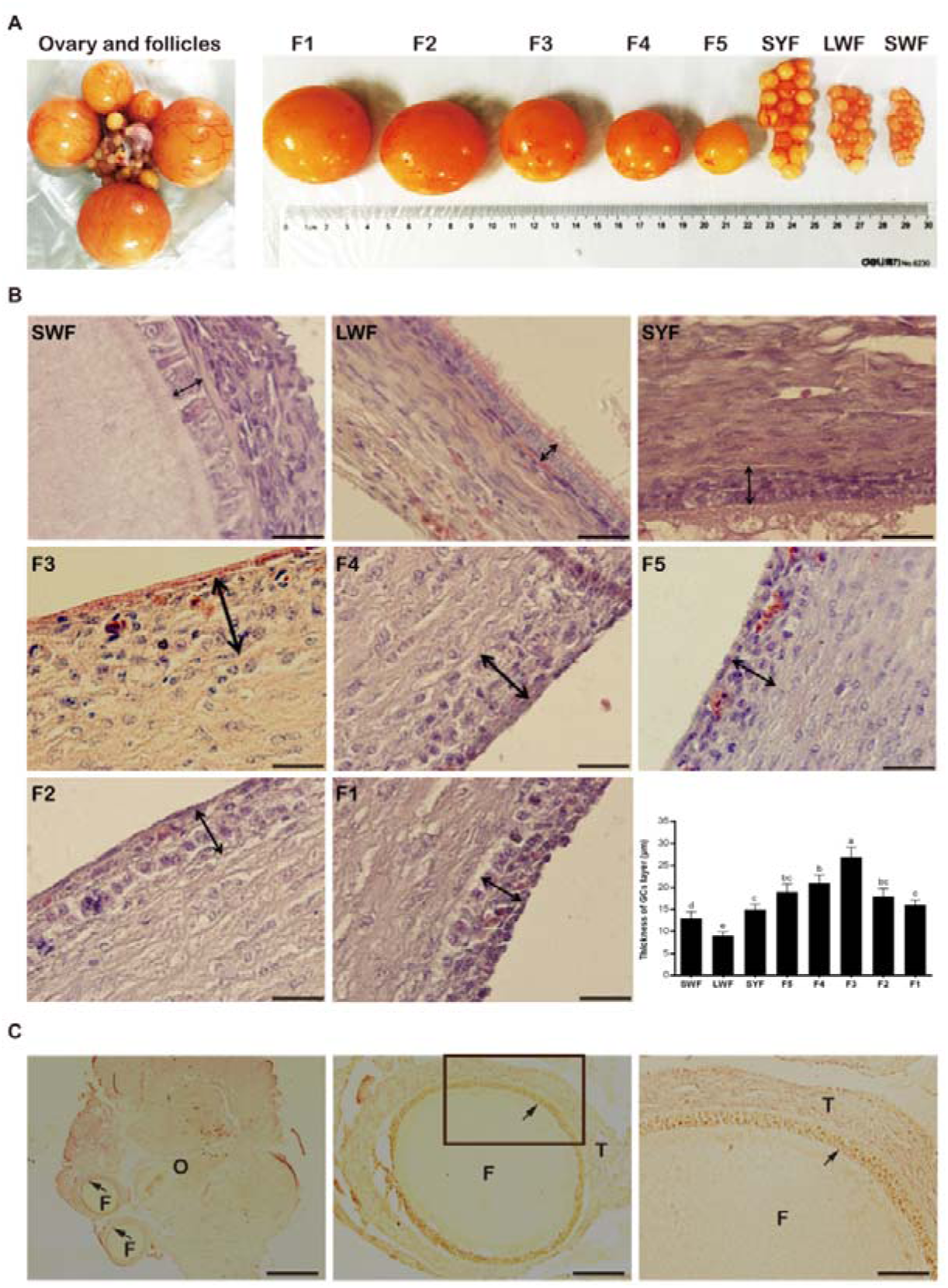
Histological analysis of goose ovary and follicle in laying period and localization of Smad9 in goose ovary. (A) Ovary, hierarchical follicles and its size. (B) Variation and comparison of the thickness of GCs layer during follicular development; *n*=3. (C) Smad9 expression in the GCs layer of goose follicle. The letter F, O and T represents the follicle, the ovary and the theca interna respectively. Arrows indicate Smad9 signaling in GCs layer. Bars with different lowercase letters indicate significant differences (*P*< 0.05). Scale bars: 20 µm in B; 500 µm in the left of C; 100 µm in the middle of C; 50 µm in the right of C.

### Smad9 activation can be transferred to the nucleus in gGCs

Smad9 is a transcription factor which must be present in the nucleus and play its biological functions. In order to locate total Smad9 and phosphorylated Smad9 (p-Smad9, the active form of Smad9 protein) expression in gGCs, immunofluorescence techniques are used to determine their distribution. The results showed that Smad9 protein is only present in the cytoplasm and p-Smad9 protein is present in the cytoplasm and nucleus (Fig. 2A,B). These results also mean that Smad9 can be translocated to the nucleus to play its biological role after activation.

**Fig. 2.**
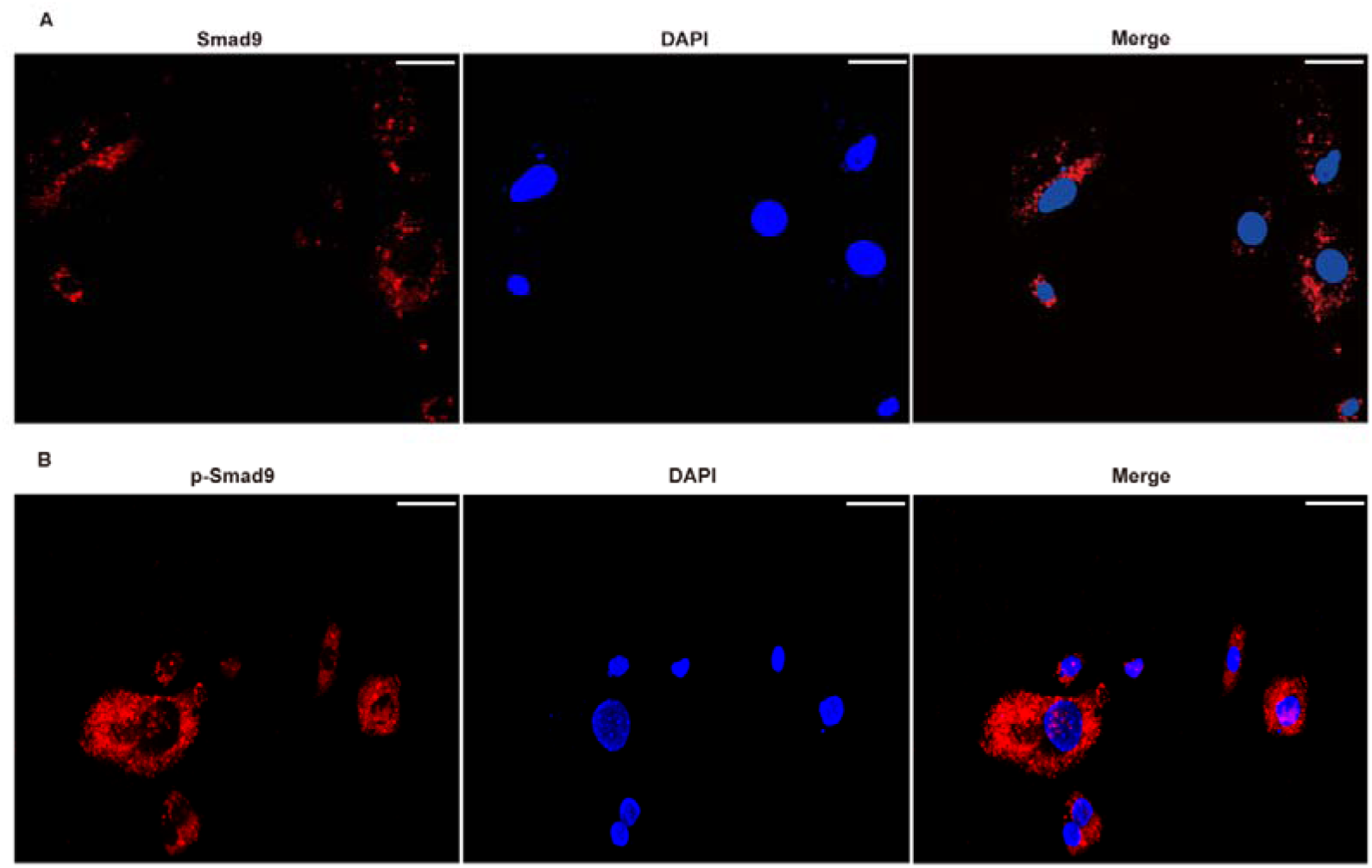
Localization of Smad9 and p-Smad9 proteins in gGCs by immunofluorescence. Cells were fixed and stained with the nuclear dye 4,6-diamidino-2-phenylindole (DAPI; blue), red represents Smad9 or p-Smad9 signaling. (A) Smad9 signaling. (B) p-Smad9 signaling. Scale bars: 50 µm.

### Differential expression of Smad9 in hierarchical follicles

To test whether the expression of Smad9 changes with the process of follicle development, we further analyzed its expression in follicles at different stages by quantitative real-time polymerase chain reaction (qRT-PCR) method. The results showed that in control group, Smad9 expression is gradually decreased in the prehierarchical follicles, which tended to decrease in the preovulatory follicles but with a fluctuation in the F4. It expression touched the bottom in the F1 follicle (Fig. 3A). In contrast, Smad9 expression is inhibited in LDN-193189 treated group, especially in the preovulatory phase with an exception of SYF (Fig. 3A,B). Moreover, the differences of Smad9 expression among SWF, LWF, SYF, F4 and F1 were significant while they were very significant among F5, F3 and F2. In addition, Smad9 transcription and translation levels were consistent in respective group, the same as to total Smad9 and p-Smad9 (Fig. 3C). In a word, these results revealed that Smad9 is differentially expressed along with the process of follicle development.

**Fig. 3.**
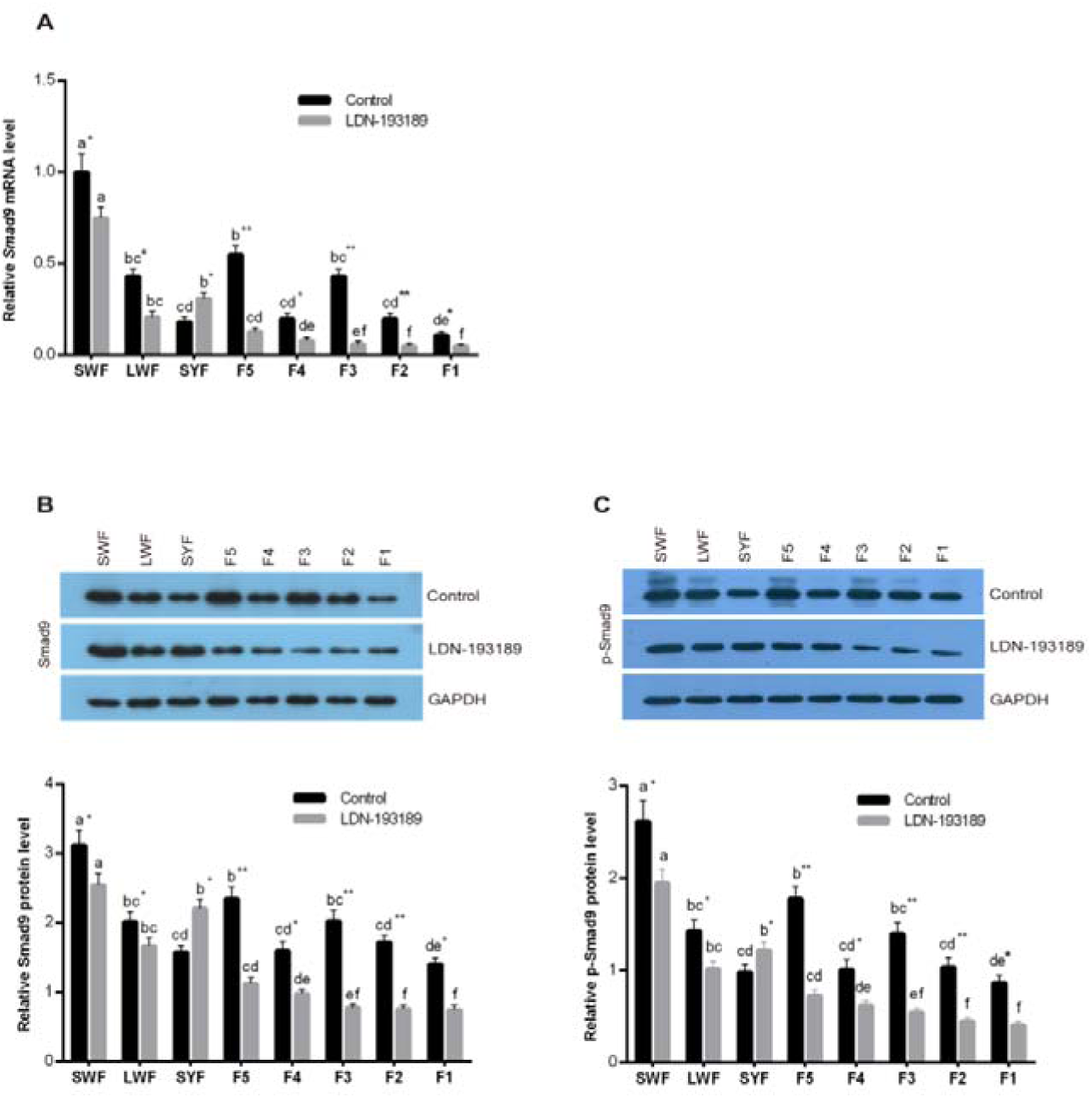
Smad9 expression in LDN-193189 treatment and control groups *in vivo*. (A) qRT-PCR analysis of Smad9 mRNA expression in LDN-193189 treatment and control groups. (B,C) Western blot analysis of Smad9 and p-Smad9 protein expression in LDN-193189 treatment and control groups. Bars with different lowercase letters are significantly different between the same group from different hierarchy follicles (*P*< 0.05). * and ** indicate significant differences between control group and LDN-193189 group in the same hierarchy follicles. * *P*< 0.05; ** *P*< 0.01; *n*=3.

### Blockage of Smad9 promotes the LHR expression and E_2_ biosynthesis as well as egg production *in vivo*

FSHR and LHR were reported to be involved in follicular development and ovulation (Camp et al., 1991; Ji et al., 2014). To investigate whether LDN-193189 treatment has an effect on the expression of these two hormone receptors, we first analyzed their expression in hierarchical follicles from laying geese by qRT-PCR method (Fig. 4A,B). The results showed that FSHR expression were different with the process of follicle development, which was not clearly affected by LDN-193189 treatment (Fig. 4A). The results also showed that LHR expression gradually increases in prehierarchical follicles until SYF, in which it was significantly higher than in other follicles (*P*< 0.05), then decreased in F5 but increased in F4, followed by a gradual decline from F3 to F1. In contrast, in LDN-193189 treatment groups, the LHR expression in prehierarchy follicles was similar in control group, however, it was different in preovulatory follicles because it gradually increased. There is no significant difference between SYF and F1, but it was significantly higher than that in any other follicles. In both the control and LDN-193189 treatment groups, the differences were not significant in SWF and LWF, and significant among F5, F3 and F2 (*P*< 0.05), and very significant among SYF, F4 and F1 (*P*< 0.01) (Fig. 4B).

**Fig. 4.**
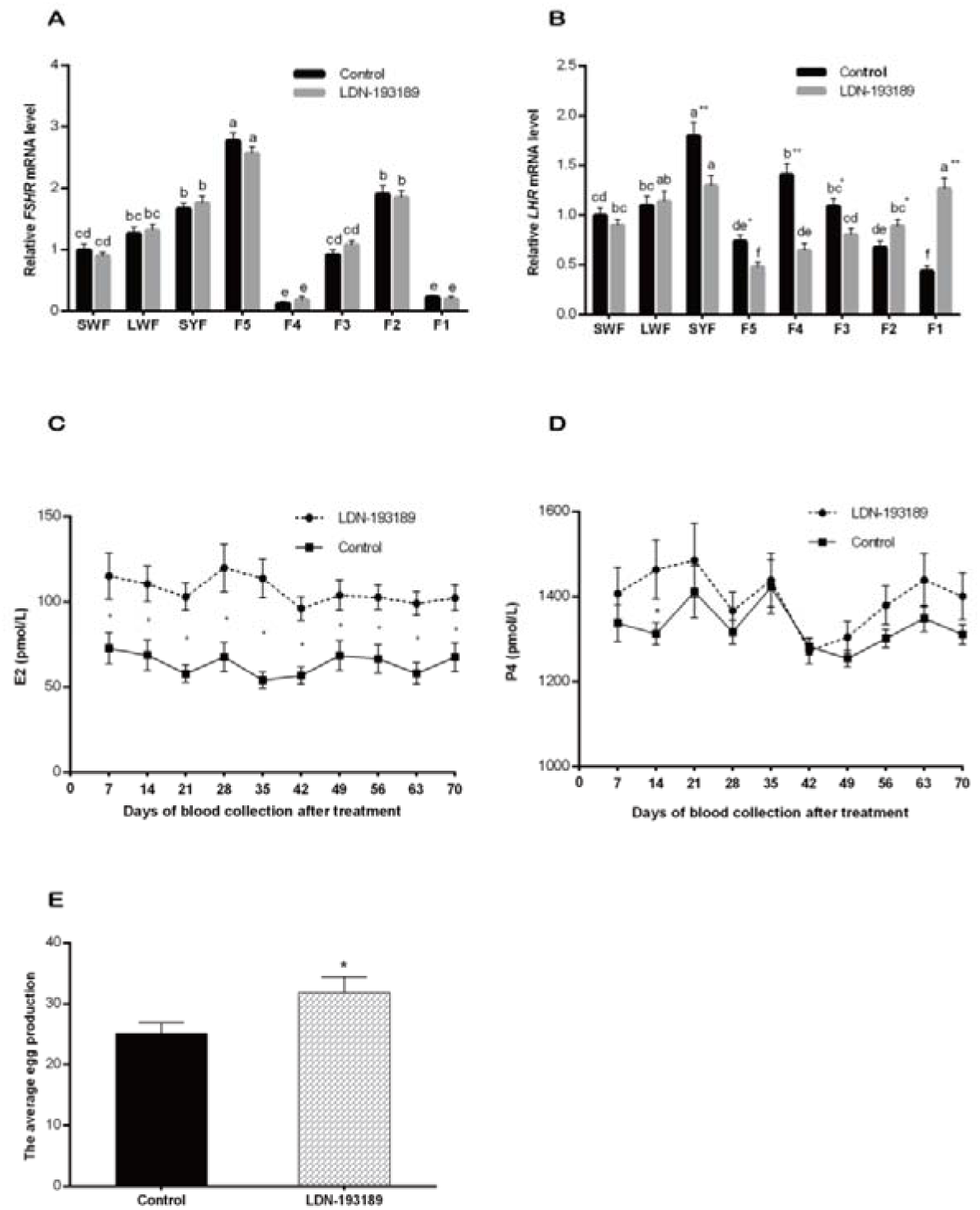
FSHR and LHR expression, E_2_ and P_4_ secretion as well as the average egg production from LDN-193189 treatment and control groups *in vivo*. (A) qRT-PCR analysis of FSHR expression in LDN-193189 treatment and control groups. (B) qRT-PCR analysis of LHR expression in LDN-193189 treatment and control groups. (C,D) Determination of E_2_ and P_4_ expression level by ELISA in control and LDN-193189 groups. (E) The average egg production in control and LDN-193189 groups (*n*=10 each). Bars with different lowercase letters are significantly different between the same group from different hierarchy follicles (*P*< 0.05). * and ** indicate significant differences between control group and LDN-193189 group in the same hierarchy follicles. * *P*< 0.05; ** *P*< 0.01; *n*=3.

E_2_ and P_4_ were two representative steroid hormones that regulate follicular development and ovulation. To study whether the treatment of LDN-193189 has any effects on the expression of these two steroid hormones, the E_2_ and P_4_ in the serum samples from laying period geese were measured by ELISA. The results showed that E_2_ levels in LDN-193189 treatment group were significantly higher than that in control group (*P*< 0.05) (Fig. 4C). However, P_4_ levels were significantly higher in the treatment group than that in the control group only in 14 days post LDN-193189 treatment (*P*< 0.05). Although there were no significant differences were found at other points of time, a slight increase was seen after treatment with LDN-193189 (Fig. 4D). Most importantly, the average egg production of LDN-193189 treatment group (31.86 eggs) was significantly higher than that of control group (24.78 eggs) (*P*< 0.05) (Fig. 4E). In conclusion, blockage of Smad9 *in vivo* significantly affects E_2_ secretion in serum and folliclar LHR gene expression which is closely related to follicular development and ovulation, eventually contributing to promoting egg production.

### Blockage of Smad9 promotes the LHR expression and E_2_ biosynthesis and cell proliferation of GCs cultured *in vitro*

GCs play a vital role in the development and maturation of follicles (Tilly et al., 1991). To further elucidate the effects of Smad9 on the development of GCs, we examined the relationship between Smad9 expression and hormone synthesis, hormone receptor expression and cell proliferation after primary cultured gGCs being treated with BMP-4, BMP-4 / LDN-193189, Smad9-siRNA for 48 h respectively (Fig. 5). Smad9 mRNA and protein were detected by qRT-PCR and western blotting, respectively. The results showed that Smad9 gene was knocked down by Smad9-siRNA (Fig. S1). Smad9 expression was significantly higher in BMP-4 group than that in blank (*P*< 0.05), and that in Smad9-siRNA group and BMP-4 / LDN-193189 group were significantly lower than that in blank (*P*< 0.05). The expression of p-Smad9 was similar to the expression of total Smad9 in those groups (Fig. 5A,B).

**Fig. 5.**
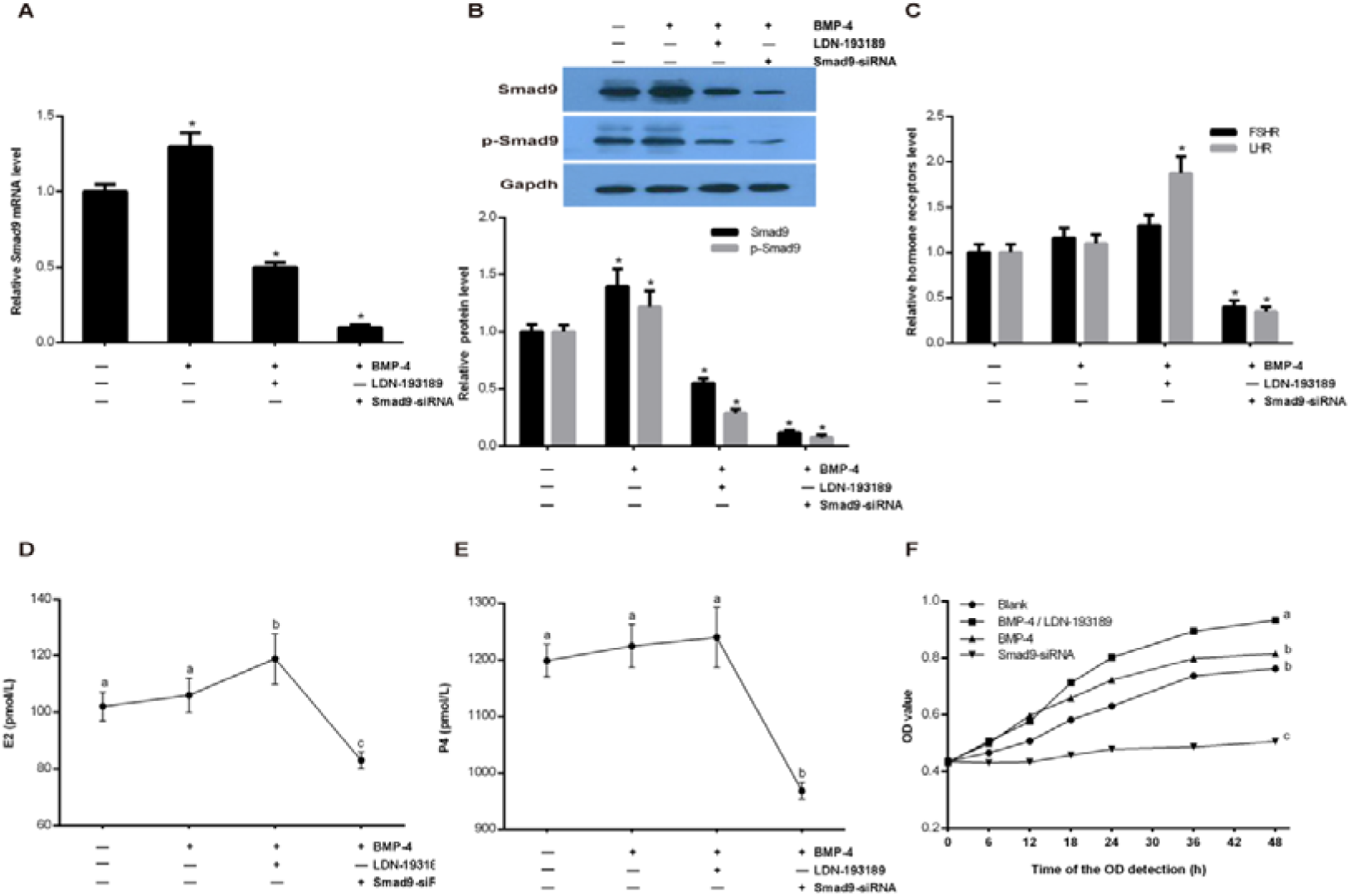
FSHR and LHR expression, E_2_ and P_4_ secretion as well as cell proliferation are related to Smad9 expression which was treated by BMP-4, BMP-4 / LDN-193189 and Smad9-siRNA respectively for 48 h in cultured gGCs. (A) qRT-PCR analysis of Smad9 expression. (B) Western blot analysis of Smad9 and p-Smad9 proteins expression. (C) qRT-PCR analysis of FSHR and LHR expression. (D,E) ELISA measurement of E_2_ (D) and P_4_ (E) levels. (F) Analyses of GCs growth using CCK-8 assay. Bars with different lowercase letters are significantly different between groups (*P*< 0.05). * represents a significant difference compared to the blank group. * *P*< 0.05; *n*=3.

The qRT-PCR analysis showed that FSHR expression were no significant difference in BMP-4 group and BMP-4 / LDN-193189 group compared with blank, however, there was significantly lower in Smad9-siRNA group than in blank (*P*< 0.05). LHR expression in BMP-4 / LDN-193189 was significantly higher than in the blank (*P*< 0.05), and there was no significant difference in BMP-4 compared with blank. However, it was significantly lower in Smad9-siRNA than that in blank (*P*< 0.05) (Fig. 5C).

E_2_ and P_4_ in the supernatants were analyzed by ELISA showed that E_2_ levels were significantly higher in BMP-4 / LDN-193189 than that in the blank (*P*< 0.05), no significant difference in BMP-4 group compared with the blank. P_4_ levels in BMP-4 group and BMP-4 / LDN-193189 group were no significant difference in comparison with the blank. However, the concentration of E_2_ and P_4_ were significantly lower in Smad9-siRNA than in the blank (*P*< 0.05) (Fig. 5D, E).

The growth of GCs was examined by CCK-8 assay. The results showed that BMP-4 / LDN-193189 markedly prompted the growth of GCs compared with the blank (*P*< 0.05). And we also found that Smad9-siRNA markedly inhibited the growth of GCs compared with the blank (*P*< 0.05). In addition, there was a slight increase in promoting cell proliferation efficiency in BMP-4 group compared with blank, but no significant difference (Fig. 5F). In conclusion, Smad9 plays an important role in the development of GCs by affecting the secretion of E_2_ and LHR expression, which can promote cell proliferation.

### Smad9 regulates LHR transcription in gGCs

Unexpectedly, FSHR *in vivo* and *in vitro* experiments showed no significant difference, which promoted us to test whether Smad9 binds to LHR promoter by ChIP assay. Our results showed that Smad9 can bind to the promoter region of LHR and regulating its transcription (Fig. 6A). It was remarkable that Smad9 expression increased post-stimulating with BMP-4 in gGCs and also increased its combined efficiency with LHR promoter (Fig. 6B). Intriguingly, the efficiency of Smad9 bonding to the LHR promoter was not reduced because Smad9 phosphorylation was inhibited, meanwhile, LHR transcription increase was observed after inhibiting Smad9 protein phosphorylation with LDN-193189 (Fig. 6B,C). In conclusion, this inner molecular mechanism shows us that Smad9 can directly regulate LHR transcription in gGCs. In addition, the proposed pathway of Smad9-mediated LHR transcription is summarized in Figure 7.

**Fig. 6.**
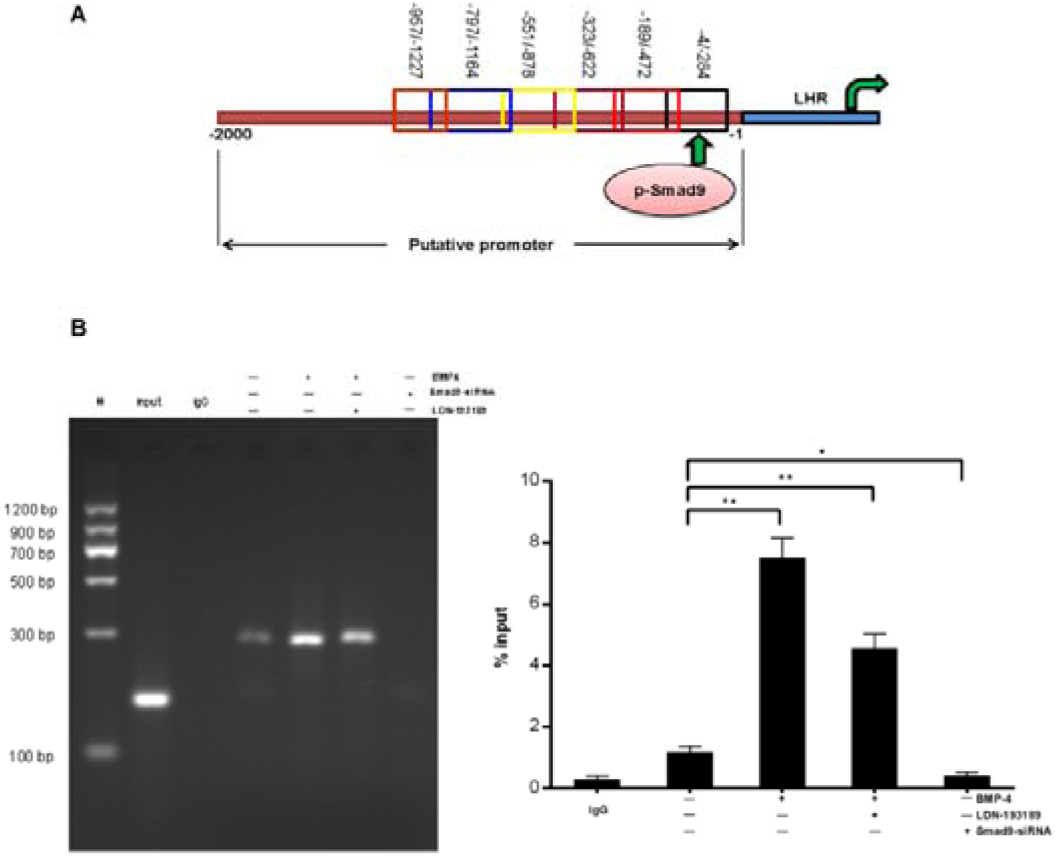
Smad9 directly regulates LHR transcription in gGCs. (A) The cartoon shows the existence of Smad9 protein binding site at the upstream region of the LHR promoter. (B) The cell lysates from gGCs samples pull-down experiments showed different levels of enrichment in indicated groups. * *P*< 0.05; ** *P*< 0.01; *n*=3.

**Fig. 7.**
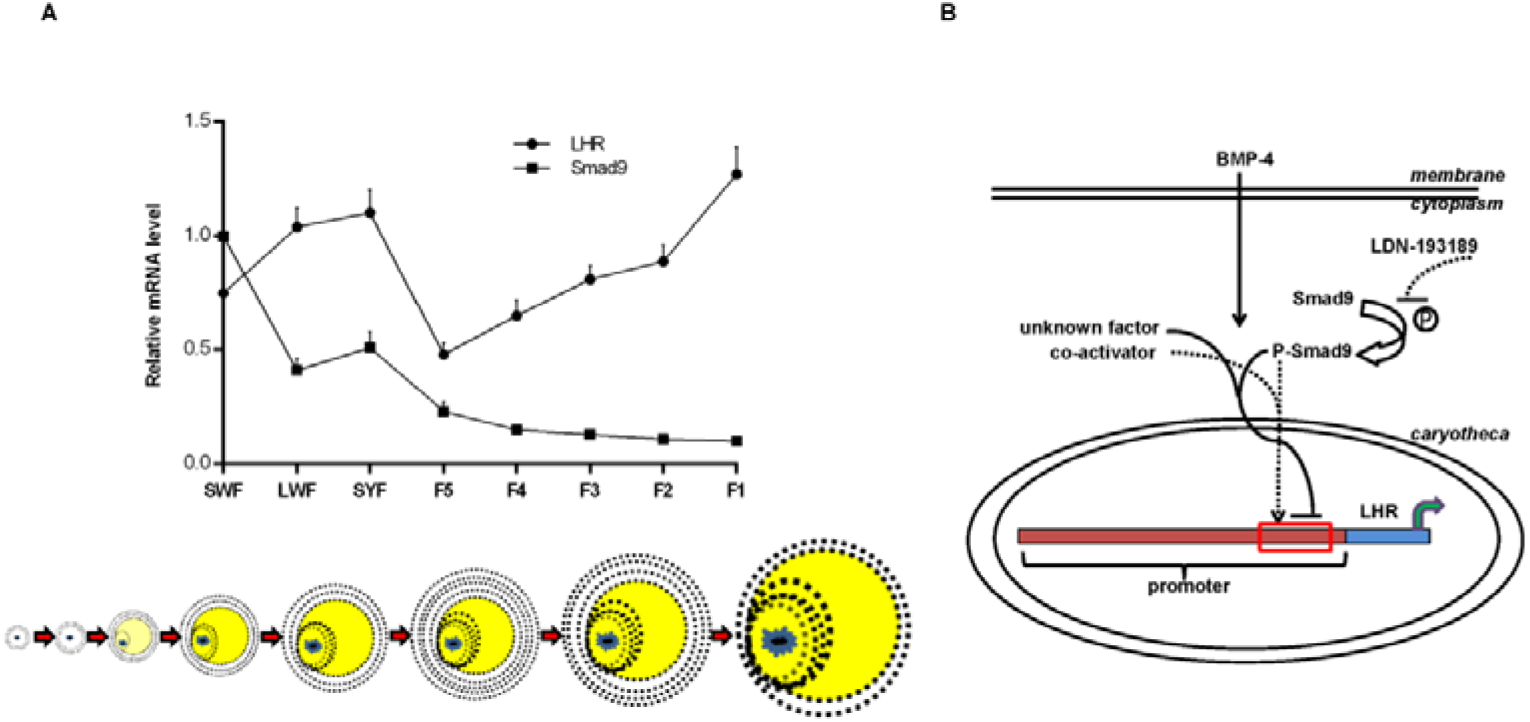
The possible patterns and mechanisms of Smad9 regulating follicular development and ovulation in goose. (A) Smad9 expression and LHR transcription synergistic model with the development of follicles when Smad9 is suppressed. (B) One mechanism is conjectured by which Smad9, served as a sensor of balance, regulated LHR transcription. The solid line represents the control mode of the high expression of Smad9 and the dashed line represents the control mode when Smad9 is suppressed.

## DISCUSSION

To our best knowledge, the present study is the first to characterize the expression and localization of Smad9 in goose hierarchical follicles and follicular GCs during the laying stage. Our results clearly showed that Smad9 is majorly located in the cytoplasm while the p-Smad9 is translocated to the nucleus as was described in the previous study (Massague, 2012). Also, for the first time our results revealed that appropriate level of Smad9 expression in gGCs during reproductive cycle is positively correlated with E_2_ secretion, LHR transcription and cell proliferation. In addition, our H&E staining results indicated that the thickness of the GCs layer changed with follicular development, which further showed the importance of GCs to follicular development.

### Particularity of Smad9 distribution

GCs play an essential role in the development, maturation and ovulation of follicle (Fitzharris and Baltz, 2006; Makita and Miyano, 2014; Orisaka et al., 2009; Zhang et al., 2011). It has previously been indicated that interrupted endogenous BMP/Smads signaling influence the growth and steroidogenesis of porcine GCs (Wang et al., 2010). Smad9 is an important downstream transcription factors of BMP/Smads signaling pathway which was ubiquitously expressed in different tissues and mediates various physiological processes (Massague and Wotton, 2000). In our study, the changing localization pattern of Smad9 in developing follicles GCs indicated the implication of BMP/Smads signaling in goose ovary. Accordingly, we may infer that similar to other mammals, BMP/Smads system in regulating female reproduction also occur in geese. We can also infer that the presence of Smad9 in GCs to facilitate the reception and transduction of signals inside and outside of the follicle.

### Smad9 difference expression in the hierarchical follicles of goose

In this study, hierarchy follicles with continuous size have been successfully obtained from laying goose. Further studies demonstrated that the morphology and quantity of the follicles used in our study were similar to that from other birds (Etches and Petitte, 1990). We separated the follicular GCs and detected the expression of Smad9. For the first time the spatiotemporal patterns of Smad9 expression in the hierarchical follicles of goose were revealed. Our data showed that Smad9 is differentially expressed in different hierarchical follicles at both transcription and translation levels. Also, our present results further extended our previous findings on the expression of Smad9 in goose ovary (Xu et al., 2015). The differential expression of Smad9 during follicular hierarchy establishment indicates its involvement in establishing or maintaining the follicular hierarchy and regulating follicle development.

### Smad9 and follicular development and ovulation

It is well known that the more the follicles that been chosen to enter the hierarchy, the longer the egg laying period and higher production of poultry. The development of avian follicles is a complex process, which is regulated by different hormones and signaling from various receptors such as E_2_ and P_4_, FSHR and LHR (Caicedo Rivas et al., 2016; Calvo and Bahr, 1983; Ji et al., 2014; Jia et al., 2010; McGee and Hsueh, 2000; Qin et al., 2013; Regan et al., 2015; Wei et al., 2013). In this study, the BMP type|kinase inhibitor, LDN-193189, was used to regulate Smad9 expression *in vivo* (Cannon et al., 2010; Derwall et al., 2012; Lee et al., 2011; Mayeur et al., 2015; Yu et al., 2008). We found that post treatment of LDN-193189, Smad9 expression was downregulated in hierarchical follicles, especially in the preovulatory phase. Meanwhile, we also found that the E_2_ and LHR levels were significantly increased, the P_4_ and FSHR levels were not changed. The results from our *in vitro* experiment further verified the reliability of results from *in vivo*. Our data also indicated that LHR expression in LDN-193189 treatment group from *in vivo* was consistent with high yield poultry chickens (Liu and Zhang, 2008). Intriguingly, we were surprised to find that the average egg production was significantly increased in LDN-193189 treatment geese compared with that in the control group (*P*< 0.05). In addition, we also found the efficiency of cells proliferation in BMP-4 / LDN-193189 group from *in vitro* was significantly higher than that in the blank group (*P*< 0.05). Both *in vivo* and *in vitro* results indicated that the expression of regulating Smad9 can directly affect E_2_ secretion, LHR transcription and cell proliferation, which finally contributes to the increased egg production. Unexpectedly, the expression of FSHR, LHR, E_2_ and P_4_ as well as GCs growth were all significantly decreased after Smad9 knock down. Our data also suggested that Smad9 inhibition may indirectly affect the other unknown mechanisms, which changes the expression of FSHR and LHR, E_2_ and P_4_ secretion and cell proliferation, remain to be further studied. Thus, our data strongly suggests that inhibition of Smad9 is a promising strategy with great potential for elevating goose egg production in the future.

### Smad9 and LHR transcription

The transition from prehierarchical follicles to preovulatory follicles is accompanied by a shift from FSHR dominance to LHR dominance in follicular development (Johnson and Woods, 2009). Therefore, LHR is critical to the ovulation of mature follicle. Our ChIP assay results showed that Smad9 can directly bind to the genomic promoter region of LHR and regulate its transcription. Although Smad9 binding efficiency was higher in BMP-4 group than that in BMP-4 / LDN-193189 treatment group, the expression of LHR was lower in BMP-4 group at mRNA level. Serving as a sensor of LHR transcription, Smad9 can keep the balance of LHR transcription to some extent, and its high or low level doesn’t work in follicular maturation. A recent report also showed that when Smad9 is expressed at high level, it could suppress BMP activity without inhibiting the phosphorylation of R-Smad (Tsukamoto et al., 2014). Taken together, these data indicate that Smad9 is the key regulator of LHR transcription in laying goose.

### Conclusions

The present study demonstrates that: (1) Smad9 is expressed in GCs of goose follicles; (2) Smad9 is differentially expressed in goose hierarchical follicles; (3) Smad9 regulates E_2_ secretion and cell proliferation; (4) Smad9 can bind to the promoter region of LHR and regulate its transcription, which is served as a sensor of balance.

## MATERIALS AND METHODS

### Animals, tissue harvest and serum samples collection

Twenty-four laying period female Chinese Wanxi White Geese (from Anhui Sanyuan Breeding Co., Ltd.) were used in this study. The geese were divided into two groups and were treated with LDN-193189 (0.15 mg-1 kg/day) via wing vein (n = 12) or with control saline (n = 12) once a day for 10 successive days. Ten sequential blood samples were obtained at weekly interval a week after treatment. The blood samples were centrifuged at 1,500 rpm for 20 min at 4℃ and the serum samples were restored at −80℃ until use. All animals had free access to water and feed. The egg production was recorded during the geese laying period. To obtain follicles from the mature ovaries, geese were sacrificed after ovipositionin the middle of a laying stage. Hierarchical follicles were collected and stored in PBS or liquid nitrogen for further experiments. The RNA and protein were isolated from the follicles after removing the yolk. All animal experiments were approved by the Institutional Animal Care and Use Ethics Committee of Anhui Normal University and performed in accordance with the “Guidelines for Experimental Animals” of the Ministry of Science and Technology (Beijing, China).

### Histological analysis, immunohistochemistry and immunofluorescence

For histological analysis, briefly, hierarchical follicles were fixed in bouin’s solution for 48 h and embedded with paraffin. Sections were sliced at 5 µm thickness and each section was stained with H&E (Boster, China) as described previously (Lam, 1997).

For immunohistochemistry, follicles were embedded in paraffin following standard protocols. The transverse sections (5 µm) were incubated with antibody against Smad9 (1:1000; Abcam, USA) for overnight at 4℃. The sections were then incubated with horseradish peroxidase-conjugated secondary antibodies followed by diaminobenzidine and counterstained with hematoxylin.

For immunofluorescence, GCs were grown in glass chamber slides. After treatments, cells were washed with PBS three times and fixed in 4% paraformaldehyde. For immunostaining, cells were incubated with the antibodies against Smad9 (1:1000; Abcam, USA) and p-Smad9 (1:800; CST, USA) for overnight at 4℃. Slides were washed three times with cold PBS and incubated with donkey anti-rabbit IgG-FITC (1:1000; Beyotime, China) for 1 h at RT. Then, the slides were washed three times with cold PBS and cells were counterstained with DAPI (Beyotime, China). The immunofluorescent signals were examined with Olympus BX61 fluorescence microscope. All images were captured using the same settings and saved in the same format.

### GCs culture, design and transfection of Smad9-siRNA and supernatant Samples collection

The GCs of SYF follicle were separated using the improved method as described by Gilbert et al (Gilbert et al., 1977). Then the cells were incubated at 39℃ in a water-saturated atmosphere of 95% air and 5% CO2.

Six oligonucleotides were designed and synthesized by Shanghai GenePharma Co., Ltd according to the cDNA sequence of goose Smad9 (Table S1). The specificity of these sequences was then confirmed by transfection preliminary experiment according to the manufacturer’s instructions (GenePharma, China). Finally, the siRNA sequence of Smad9 was identified as siRNA-1.

The cultured GCs were randomly divided into five groups: blank, BMP-4 50 ng/ml, BMP-4 50 ng/ml / LDN-193189 100 nM, Smad9-siRNA 16.7 nM and NC-siRNA 16.7 nM. Cells and culture supernatants were collected separately after treatment with serum-free medium for 48 h. The cells were collected RNA and protein preparation, and supernatant were restored at −80℃ until use. Three replicates for each group and three independent repeated for each experiment.

### Cell proliferation assay

A total of 3 × 10^3^ GCs per well were incubated for 24 h in a 96-well plate. Cell proliferation was determined by using cell counting kit-8 (CCK-8) assay kit (Beyotime, China) after treatment with BMP-4, BMP-4 / LDN-193189, Smad9-siRNA and blank for 6 h, 12 h, 18 h, 24 h, 36 h and 48 h following the instruction from the manufacturer. The experiment was repeated for three times.

### Quantitative real-time PCR

Total RNA was extracted from follicles or cultured GCs using Trizol (Invitrogen, USA) according to the manufacturer’s instructions. The concentration and purity of isolated total RNA were determined using a standard spectrophotometer (K.O., China). The cDNA was obtained using reverse transcription kit (Tiangen, China) according to the manufacturer’s instructions. The qRT-PCR was performed in a Bio-Rad CFX Manager System (BIO-RAD, USA) using a qRT-PCR kit and SYBR Green as the detection dye (Tiangen, China) according to the manufacturer’s instructions. Primers used for qRT-PCR are listed in Table S2. Each sample was repeated three times, and the relative concentrations of all interested genes were calculated using the 2^−△△^Ct^^ method (Livak and Schmittgen, 2001).

### Western blotting

Total proteins were extracted from goose follicle GCs or cultured GCs with RIPA lysis buffer (Beyotime, China) and concentration was quantified using bicinchoninic acid (BCA) protein assay kit (Huamei, China) according to the manufacturer’s instructions. Protein extracts were subjected to western blot analysis with rabbit anti-p-Smad9 antibody (1:1000; CST, USA), mouse anti-Smad9 antibody (1:1600; Abcam, USA), mouse anti-Gapdh antibody (1:500; Boshide, China) for primary antibodies, and anti-rabbit and anti-mouse IgG-HRP for secondary antibody, respectively.

### ELISA

The levels of E_2_ and P_4_ in serum and supernatant samples were measured by goose E_2_ and P_4_ ELISA kit (Halin, China) according to the manufacturer’s instructions, respectively.

### ChIP assay

Three dishes of GCs were used, one dish as blank, another two dishes were induced with BMP-4, one of which was stimulated by LDN-193189 after BMP-4 inducing for 12 h, and then cultured for 48 h. ChIP was conducted using a ChIP assay kit (CST, USA) following manufacturer’s instructions. GCs were cross-linked with 1% formaldehyde (Sangon, China). Chromatin was digested with micrococcal nuclease (CST, USA). The immune complexes of DNA-protein were immune precipitated by p-Smad9 antibody (CST, USA). Normal rabbit IgG (CST, USA) was used as control. The purified DNA used as a template to expand the promoter region of the LHR by qRT-PCR. Six pairs of primers were designed in −1/−2000 bp of the LHR promoter region and listed in Table S3.

### Statistical analysis

Two-tailed Student’s *t*-test for two samples was used to calculate the *p*-values. The results were presented as mean ± s.e.m, and asterisks or different letters in figures represents the level of significance as follows: “*” or “a, b, c..” *P*< 0.05 were considered statistically significant, ** *P*< 0.01 were considered statistically very significant.

## Acknowledgements

We thank Anhui Sanyuan Breeding Co., Ltd. for providing the experimental geese. We also thank Dr Yongguang Yang for critical reading of the manuscript.

## Author contributions

Y.F.C., D.L.Y. and Jun L. designed the experiments. D.L.Y., L.Z., H.J.W., Z.M.Z., Jie L., C.F.X. and F.H.C. performed the experiments. D.L.Y., J.C. and H.L.L. analyzed the data. D.L.Y. and Y.F.C wrote the manuscript.

## Data availability

Goose Smad9, FSHR, LHR and Gapdh gene data are available in NCBI with accession number XM_013192591, XM_013192472, XM_013192443 and XM_013199522, respectively.

